# Captivity and exposure to the emerging fungal pathogen *Batrachochytrium salamandrivorans* are linked to perturbation and dysbiosis of the amphibian skin microbiome

**DOI:** 10.1101/339853

**Authors:** Kieran A. Bates, Jennifer M.G. Shelton, Victoria L. Mercier, Kevin P. Hopkins, Xavier A. Harrison, Silviu O. Petrovan, Matthew C. Fisher

## Abstract

1. The emerging fungal pathogen, *Batrachochytrium salamandrivorans* (*Bsal*) is responsible for the catastrophic decline of European salamanders and poses a threat to amphibians globally.
2. The amphibian skin microbiome is strongly associated with disease outcome for several host-pathogen systems, yet its role in *Bsal* infection remains unresolved. In addition, many *in-vivo Bsal* studies to date have relied on specimens that have been kept in captivity for long periods without considering the influence of environment on the microbiome and how this may impact the host response to pathogen exposure.
3. We characterised the impact of captivity and exposure to *Bsal* on the skin bacterial and fungal communities of two co-occurring European newt species, the smooth newt (*Lissotriton vulgaris*) and the great-crested newt (*Triturus cristatus*).
4. *Bsal* infection and subsequent mortality in both newt species was associated with perturbation of the skin microbiome and possible dysbiosis. In addition, reduced microbial diversity and changes in microbiome structure accompanied the transition of newts from the wild to captivity, suggesting a possible decline in microbe-associated protection and increased risk of infection by opportunistic pathogens.
5. Our findings advance current understanding of the role of host-associated microbiota in *Bsal* infection and highlight important considerations for *ex-situ* amphibian conservation programmes.

## Introduction

Microbial communities associated with amphibian skin are increasingly recognised for their ecological complexity and importance in pathogen defence. In particular, studies have demonstrated that skin-associated bacteria are linked to disease outcome in amphibians infected by the chytrid fungus *Batrachochytrium dendrobatidis* (*Bd*) (Lauer *et al*. 2007; Harris *et al*. 2009; Jani & Briggs 2014; Kueneman *et al*. 2016; Bates *et al*. 2018). Specifically, host-associated microbes may offer protection through production of pathogen-inhibiting compounds (Brucker *et al*. 2008; Woodhams *et al*. 2017), preventing pathogen colonisation (Buffie & Pamer 2013) or by outcompeting harmful microbial invaders (Kamada *et al*. 2012). In addition to the host benefits conferred by the microbiome, some microbes can promote pathogen growth (Stacy *et al*. 2016), while perturbations of host-associated microbial communities can negatively impact host health in a process called dysbiosis (Croswell *et al*. 2009). In recent years, the recognition of the microbiomes’ role in disease has led to a search for pathogen-inhibiting probiotics and microbial manipulations that could mitigate host infection and subsequently be utilised as a tool in wildlife conservation (Bletz *et al*. 2013; Kueneman *et al*. 2016).

In 2013 a novel pathogenic chytrid fungus, *Batrachochytrium salamandrivorans* (*Bsal*), was discovered that has caused mass mortalities of caudates in Europe (Martel *et al*. 2013) and threatens amphibians worldwide (Yap *et al*. 2015). While *Bd* and *Bsal* are closely related phylogenetically and occupy similar niches as the only known species within the Chytridiomycota capable of infecting vertebrates (Berger *et al*. 1998; Martel *et al*. 2013), they show marked differences in their biology. *Bd* infects over 500 amphibian species (Fisher, Garner & Walker 2009) that span all amphibian orders, whereas *Bsal* has a narrower host range limited mostly to caudates (Martel *et al*. 2014). *Bd* and *Bsal* also differ in their pathogenesis with *Bd* causing hyperkeratosis and hyperplasia of the amphibian epidermis, compared to lesions and focal necrosis in *Bsal* (Martel *et al*. 2013). Prior studies have shown variability in *Bsal* susceptibility between caudate species (Martel *et al*. 2014), however little is known of the determinants of disease outcome. Given its importance in *Bd* infection, the amphibian skin microbiome is a candidate driver of within- and between-species variability in response to *Bsal* exposure. While a great deal is known regarding the impact of *Bd* on the host microbiome, no *in-vivo* studies have investigated the microbiome response to *Bsal*. Importantly, it is not possible to predict the impact of *Bsal* on amphibian microbiota based on prior *Bd* studies, or to presume a microbiome response similar to that of *Bd*. This is due to a range of factors including intrinsic biological differences between *Bd* and *Bsal* (Farrer *et al*. 2017), and the variable responses that single bacterial strains can have with different pathogen isolates. For example, it is well established that the same bacterial strain can be either inhibitory or growth-promoting depending on what genotype of *Bd* it is in co-culture with (Antwis *et al*. 2015; Antwis & Harrison 2018). In addition, more recent studies have shown that certain bacteria isolated from amphibian skin inhibit *Bsal*, but not always *Bd, in-vitro* (Muletz-Wolz *et al*. 2017) and that *Bd* and *Bsal* metabolites can modulate growth of different bacteria (Woodhams *et al*. 2017). Taken together, these findings show that differences exist in the way *Bsal* and *Bd* interact with the microbiome of amphibians, reinforcing the importance of investigating the *in-vivo* host response to *Bsal* exposure.

A key component of wildlife disease mitigation for highly threatened species is the establishment of assurance populations or disease treatment in captivity (Mendelson *et al*. 2006; Gascon 2007). However, despite recognition of the importance of the microbiome in host health and pathogen defence, few studies have investigated the potential impact of captivity on the amphibian skin microbiome (Becker *et al*. 2014; Loudon *et al*. 2014; Bataille *et al*. 2016; Kueneman *et al*. 2016; Sabino-Pinto *et al*. 2016) and none have investigated this with respect to *Bsal* mitigation. Prior studies have yielded mixed results with reductions in bacterial alpha diversity and depletion of chytrid-inhibiting bacteria in captive compared to wild individuals for some host species (Loudon *et al*. 2014; Bataille *et al*. 2016; Kueneman *et al*. 2016; Sabino-Pinto *et al*. 2016) while increased alpha diversity was seen in other species (Becker *et al*. 2014). In addition, studies investigating the impact of captivity on the amphibian skin microbiome have neglected to examine microbial kingdoms other than bacteria, despite recent advances demonstrating that fungi may be equally important to host health and disease resistance (Kearns *et al*. 2017). Further, cross-kingdom responses to captivity may not be uniform making it essential that a more holistic outlook of the skin microbiome is taken. Understanding the effect of captivity on the host microbiome is especially important with regard to *Bsal* exposure since field based interventions are unlikely to be successful owing to disease transmission occurring at low population density (Schmidt *et al*. 2017) and with the only currently effective treatments being captivity based (Blooi *et al*. 2015a; Blooi *et al*. 2015b). Gaining insights into how both *Bsal* exposure and the transition from the wild to captivity affect the caudate skin microbiome is therefore an important advancement in our understanding of infection as well as being valuable in informing future captivity-based conservation interventions.

In this study, we combine field and laboratory studies to investigate how the amphibian skin microbiome changes with the transition from the wild to captivity followed by subsequent exposure to *Bsal*. We focus on two UK caudate species, the smooth newt (*Lissotriton vulgaris*) and the great crested newt (*Triturus cristatus*). While *L. vulgaris* is ubiquitous in the UK, *T. cristatus* is rarer, more localised and declining in many parts of its natural range (Edgar, Griffiths & Foster 2005). *T. cristatus* is also listed as a protected species in Annexes II and IV of the European Commission Habitats Directive and under the UK Wildlife and Countryside Act 1981. Consequently, the long-term population viability of *T. cristatus* is particularly vulnerable to local disease outbreaks. The susceptibility of captive-raised *T. cristatus* to *Bsal* has been tested in a prior study (Martel *et al*. 2014) which showed mortality in all infected animals. Meanwhile, no study to date has investigated the lethality of *Bsal* in *L. vulgaris*. Testing the effect of *Bsal* on endemic UK species and the possible risk it poses to wild populations is vital given the recent emergence of *Bsal* in private collections (Cunningham *et al*. 2015) and the continued spread of pathogenic chytrids through the global trade (O’Hanlon *et al*. 2018) suggesting a wild outbreak is possible. Understanding the effects of *Bsal* exposure and the influence of captivity on the amphibian skin microbiome could therefore not only improve the capacity for developing adequate national response protocols to disease outbreaks, but also inform effective captivity based measures that seek to maximise natural pathogen protection.

## Methods

### Field sampling and captivity experiment

A total of 15 adult *Triturus cristatus* and 15 adult *Lissotriton vulgaris* were collected from a reserve in Cambridgeshire, UK. Individual newts were generally found under rocks at night and represented less than 0.1% of the total estimated site population. Using a single sterile MW100 rayon tipped dry swab (MWE Medical Wire, Corsham, UK), the skin microbiome was sampled by swabbing the ventral and dorsal surfaces 10 times, and the fore- and hindlimbs five times. Swabs were stored at −80°C until processed. Animals were transferred to individual 1.6L plastic boxes containing moss collected from the field site and transported to the Central Biomedical Services (CBS) Unit at Imperial College London. In captivity animals were housed individually under semi-natural conditions in plastic boxes containing a damp paper towel substrate and a cover object. Enclosures were cleaned with Rely+On Virkon (Antect International Ltd., Suffolk, UK) and animals were fed mealworms (*Tenebrio molitor*) or crickets (*Acheta domesticus*) *ad libitum* twice weekly. The animal room was kept on a 12 hour light/dark cycle and was maintained at 16°C. At two weeks post capture animals were swabbed again to measure the effects of captivity on the skin bacterial and fungal community.

### Bsal exposure experiment

In order to compare species response to *Bsal*, experiments were designed to be as similar as possible to those described in a previous study (Martel *et al*. 2014). *Batrachochytrium salamandrivorans* (isolated from a *Salamandra salamandra* outbreak in the Netherlands, isolate AMFP13/1) was grown in 25cm^3^ Nunc tissue culture flasks (Thermo Fisher Scientific, Massachusetts, USA) containing mTGhL liquid media (8g tryptone, 2g gelatin hydrosylate, 4g lactose, 950ml distilled water) and incubated at 15°C. Ten individuals from each caudate species were randomly assigned to a treatment group and exposed to 500uL of mTGhL media containing 50 × 10^4^ *Bsal* zoospores. The remaining five individuals from each species were assigned to a control group and exposed to 500uL of mTGhL liquid media. The inoculum was pipetted directly onto the dorsum of the animal. During exposure, *T. cristatus* were placed individually in 0.7L plastic boxes and *L. vulgaris* were placed in sterile petri dishes for 22 hours.

Animals were weighed and swabbed prior to *Bsal* infection on day 0 of the experiment and then every 7 days post infection for a period of 58 days. On day 58 of the experiment surviving animals were euthanized by an overdose of tricaine methanesulfonate (MS222) and subsequent destruction of the brain following UK Home Office animal procedure guidelines. In agreement with ethical protocols, any animals exhibiting pre-defined endpoint criteria (lack of righting reflex within five seconds of being inverted, persistent skin lesions covering over 20% of the body or that became septic, greater than 20% loss in body weight) were euthanized prior to day 58.

### Sample processing, DNA extraction and quantification of Bsal infection

Genomic DNA was extracted from swabs using a bead beating protocol (Boyle *et al*. 2004) and diluted 1/10 before undergoing subsequent PCR based analyses. Quantification of *Bsal* infection load was done using qPCR amplification following a modified published method (Boyle *et al*. 2004) that included a *Bsal* specific probe (STerCVIC), forward primer (STerFC) and reverse primer (STerT). Each sample was run in duplicate and with *Bsal* standards of 100, 10 and 1 genomic equivalents (GE). A distilled water negative control was also included. Samples were considered positive if both wells gave a GE of greater than 0.1.

### Bacterial microbiome sample processing

DNA extracted from swabs was used to amplify the V4 region of the 16S rRNA gene using custom barcoded primers and PCR conditions adapted from a prior study (Kozich *et al*. 2013). PCR conditions consisted of a denaturing step of 95°C for 15 min, followed by 28 cycles of 95°C for 20s, 50°C for 60s, 72°C for 60s and a final extension step of 72°C for 10 min. Each PCR including a negative water control was performed in triplicate. Amplicons were visualized on a 2% agarose gel and pooled yielding a final per sample volume of 24μl. Pooled amplicon DNA was purified using an Ampure XP PCR purification kit (Beckman Coulter, California, USA). Following purification, 1ul of each combined sample was pooled into a preliminary library and the concentration was determined using Qubit fluorometric quantification (Life Technologies, California, USA). Amplicon quality and incidence of primer dimer was assessed using an Agilent 2200 TapeStation system (Agilent Technologies, California, USA). A titration run of 300 sequencing cycles was performed on a MiSeq instrument (Illumina, California, USA) to quantify the number of reads yielded per sample from the preliminary library. An equimolar concentration of each sample was then pooled into a final composite library based on the index representation from the titration run and subsequently sequenced on a 500 cycle MiSeq run with a 250 bp paired-end strategy.

### Fungal mycobiome sample processing

DNA extracted from swabs was used to amplify the ITS2 region of the fungal internal transcribed spacer (ITS) using custom barcoded primers (Kozich *et al*. 2013) and the following PCR conditions: denaturing step of 95°C for 2 min, followed by 35 cycles of 95°C for 20s, 50°C for 20s, 72°C for 5 min and a final extension step of 72°C for 5 min. Each PCR plate included a negative swab control and negative water control, and was performed in duplicate. Amplicons were visualized on a 1.5% agarose gel and pooled yielding a final per sample volume of 50μl. Pooled amplicon DNA was purified using AMPure XP bead clean-up (Beckman Coulter, California, USA). Qubit fluorometric quantification (Life Technologies, California, USA) was used to determine the concentration of each purified sample, which were equimolar pooled to create the final library sample. This pooled sample was run on an Agilent 2200 TapeStation system (Agilent Technologies, California, USA) to assess amplicon distribution and presence of primer dimer. The sample underwent 300bp paired-end sequencing using v3 chemistry on an Illumina MiSeq platform.

### Bacterial microbiome analysis

Sequences were processed using MOTHUR (Schloss *et al*. 2009) following a previously described method (Kozich *et al*. 2013). Paired-end reads were split by sample and assembled into contigs. Sequences were quality filtered by removing ambiguous base calls, removing homopolymer regions longer than 8 bp, and trimming reads longer than 275 bp. Duplicate sequences were merged and aligned with 16S reference sequences from the SILVA small-subunit rRNA sequence database (Pruesse *et al*. 2007). A pre-clustering step grouped sequences differing by a maximum of 2 bp. Chimeric sequences were removed using UCHIME (Edgar *et al*. 2011) as implemented in MOTHUR. 16S rRNA gene sequences were clustered into groups according to their taxonomy at the level of order and assigned operational taxonomic units (OTUs) at a 3% dissimilarity level. Sequences were taxonomically classified with an 80% bootstrap confidence threshold using a naïve Bayesian classifier with a training set (version 9) made available through the Ribosomal Database Project (http://rdp.cme.msu.edu) (Wang *et al*. 2007). Sequences derived from chloroplasts, mitochondria, archaea, eukaryotes or unknown reads were eliminated. The number of sequences per sample ranged from 17804 to 63367. To mitigate the effects of uneven sampling (Schloss, Gevers & Westcott 2011) all samples were rarefied to 17804 sequences corresponding to the size of the lowest read sample. OTUs making up less than 0.01% of the total reads were removed (Bokulich *et al*. 2013). Downstream analysis of OTUs was carried out using the package Phyloseq (McMurdie & Holmes 2013) in R version 3.4.1 (R Development Core Team 2017).

### Fungal mycobiome analysis

Analysis of fungal communities was performed for the wild versus captive experiment only. Following sequencing, forward and reverse reads were assigned to samples according to dual index combinations and were paired using Paired-End reAd mergeR (PEAR) (Zhang *et al*. 2014). Paired-end reads were trimmed by per-base quality score using MOTHUR (Schloss *et al*. 2009) and reads shorter than 50bp or containing ambiguous base calls were removed. UCHIME (Edgar *et al*. 2011) was used to identify and remove chimeric sequences, and remaining sequences were clustered into Operational Taxonomic Units (OTUs) based on 97% similarity using Cd-hit (Li & Godzik 2006). The most abundant sequence in each OTU was used for BLASTn searches against the User-friendly Nordic ITS Ectomycorrhiza (UNITE) database (Koljalg *et al*. 2005). Unidentified sequences or those belonging to kingdoms other than “fungi” were removed, as were fungal sequences with BLASTn search result e-values >e^-20^ or identity <85%. The number of sequences per sample ranged from 3252 to 39493. Downstream analysis of OTUs was carried out using the package Phyloseq (McMurdie & Holmes 2013) in R version 3.4.1.

### Statistical analysis

To determine the effect of captivity and Bsal exposure on the microbiome, we calculated both alpha and beta diversity metrics using the phyloseq package (McMurdie & Holmes 2013) in R version 3.4.1 (R Development Core Team 2017). Shannon diversity was calculated for all samples and a mixed linear model (package lme4 (Bates *et al*. 2015)) was used to investigate changes in diversity in captive versus wild animals whilst taking into account repeated sampling. For the exposure experiment, separate mixed linear models were used for each newt species to investigate the effect of day of sampling, experimental group, mass, *Bsal* infection intensity and survival on Shannon diversity. P-values were calculated using the Kenward-Roger approximation of degrees of freedom in the afex package (Singmann *et al*. 2017). A Bray-Curtis distance matrix was used to calculate beta diversity for both the captivity study and *Bsal* challenge experiment. Beta diversity for both the captivity study and *Bsal* challenge experiment was visualised using Detrended Correspondence Analysis (DCA) plots. For the captivity study, the effects of captivity, host species and their interaction on skin microbial community structure was assessed using permutational multivariate analysis of variance (PERMANOVA) (Anderson 2001) using the adonis function in the vegan package (Oksanen *et al*. 2016). For the *Bsal* challenge experiment differences in beta diversity based on treatment, *Bsal* infection status and survival were investigated using PERMANOVA. Differentially abundant bacterial OTUs in both wild and captive individuals of each species and for each outcome group at day 28 of the infection experiment were determined using indicator analysis (Dufrene & Legendre 1997) using the labdsv package (Roberts 2016). An indicator score of ≥ 0.7 and *q*-value <0.05 was used as a cut-off (Becker *et al*. 2015; Longo & Zamudio 2017). Differences in survival among animals in the *Bsal* exposure experiment was investigated using a cox proportional-hazard regression model in the survival package (Therneau 2015) with mass at the beginning of the experiment, species, and GE at time of death or at the end of the experiment included as covariates.

## Results

Our analysis found rapid reductions in bacterial and fungal Shannon diversity associated with the transition from the wild to captivity for both host species (*p*<0.0001, Fig. 1a, b). Bacterial and fungal beta diversity differed in wild versus captive conditions (PERMANOVA, bacteria wild-captive: Pseudo-F_(1,56)_=66.84, R^2^=0.505, *p*=0.001, fungal wild-captive: Pseudo-F_(1,28)_=8.48, R^2^=0.217, *p*=0.001) and a host species effect was present for bacterial communities (PERMANOVA, host species: Pseudo-F_(1,56)_= 5.46, R^2^=0.041, *p*=0.007, host species*wild-captive: Pseudo-F_(1,56)_=4.02, R^2^=0.030, *p*=0.016, Fig. 1c-d). Interestingly, changes in alpha and beta diversity associated with captivity were mirrored for bacteria and fungi demonstrating a common response across microbial kingdoms. Captivity was associated with compositional changes in the core microbiome of both species with reductions in taxa such as *Cladosporium* and *Pseudomonas* (Fig. 1.e,f). Indicator analysis of wild versus captive specimens identified 178 and 167 differentially abundant bacterial OTUs in *T. cristatus* and *L. vulgaris* respectively (Fig. 2a,b; SI Table 1,2). Significant differences in fungal OTU abundance associated with captivity were also evident with 45 indicator OTUs for *T. cristatus* and 18 indicator OTUs for *L. vulgaris* (SI Table 3,4). Interestingly both host species demonstrated similar indicator OTU profiles of major bacterial groups (Fig. 2a,b) with changes in abundance of potentially important taxa such the Actinomycetales that are a major source of antimicrobial compounds (Berdy 2005) and *Lysobacter* (SI Table 1,2) that was previously identified as inhibitory against the closely related *Batrachochytrium dendrobatidis* (Brucker *et al*. 2008). In addition to the reduction in putatively protective microbes, in captive *T. cristatus* there was an increase in pathogenic fungi such as *Basidiobolus ranarum* (SI Table 3) which has previously been shown to cause disease in amphibians (Taylor *et al*. 1999).

**Figure 1.**
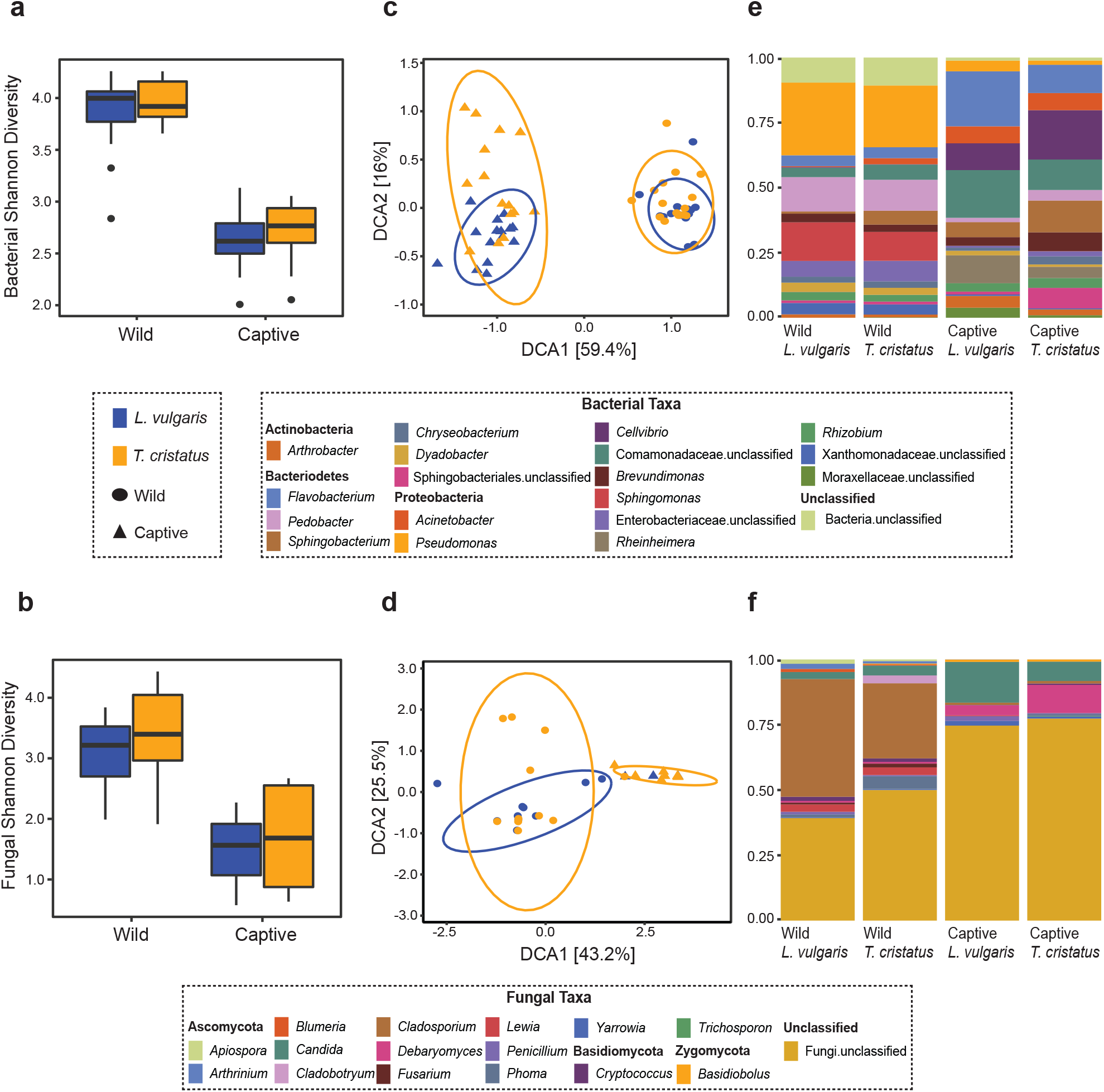
Boxplots displaying Shannon diversity in wild and captive *Lissotriton vulgaris* and *Triturus cristatus* for **(a)** bacteria **(b)** fungi. Detrended correspondence analysis (DCA) plots of beta diversity in wild and captive *L. vulgaris* and *T. cristatus* for **(c)** bacteria **(d)** fungi. Ellipses indicate 95% confidence intervals. Where ellipses are absent, insufficient samples were present. Stacked bar plots of **(d)** bacterial taxa with relative abundance > 1 % and **(e)** fungal taxa with relative abundance > 0.5%.

**Figure 2.**
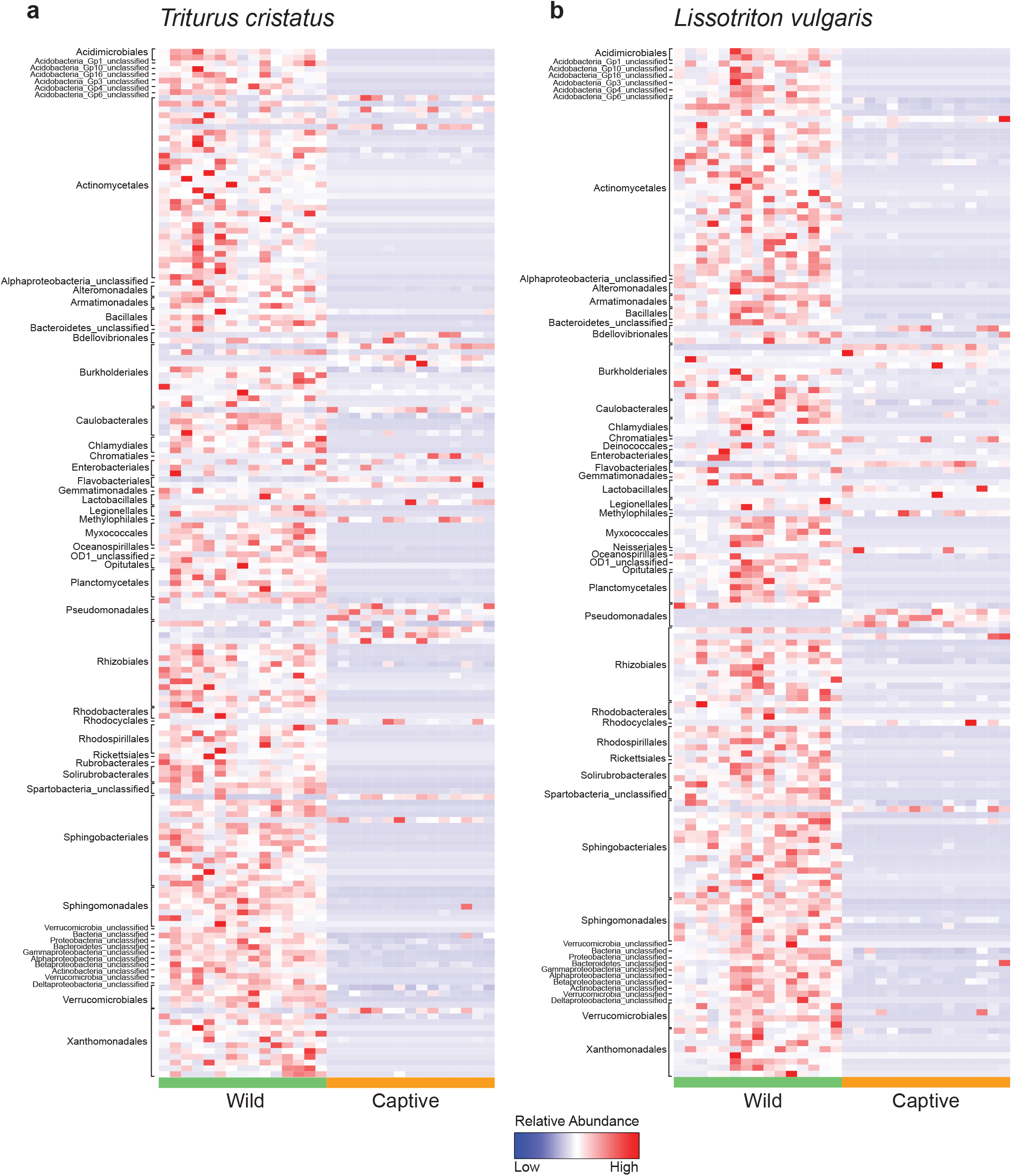
Heatmap of normalised relative abundance of bacterial indicator OTUs for wild and captive animals labelled by order for **(a)** *T. cristatus* **(b)** *L. vulgaris*.

*Bsal* exposure resulted in infection in 40% of *L. vulgaris* and 60% of *T. cristatus*. Prevalence and infection intensity fluctuated throughout the experiment for both species (SI Table 5, Fig. 3) with *T. cristatus* exhibiting consistently higher infection intensity and prevalence than *L. vulgaris*. Lesions were evident in 50% of *T. cristatus* and 75% of *L. vulgaris* that tested *Bsal* positive (SI Table 5, SI Fig. 1). Of the animals that tested positive for *Bsal*, 50% of *T. cristatus* died, while 25% of *L. vulgaris* died over the 58 days of the experiment. Of the four *L. vulgaris* that became infected with *Bsal*, three animals cleared infection, while for *T. cristatus* only one of six *Bsal* positive animals cleared infection. Survival analysis showed that infection intensity but not species or mass were significantly associated with mortality (hazard-ratio=1.07, *p*=0.022, SI Table 6).

**Figure 3.**
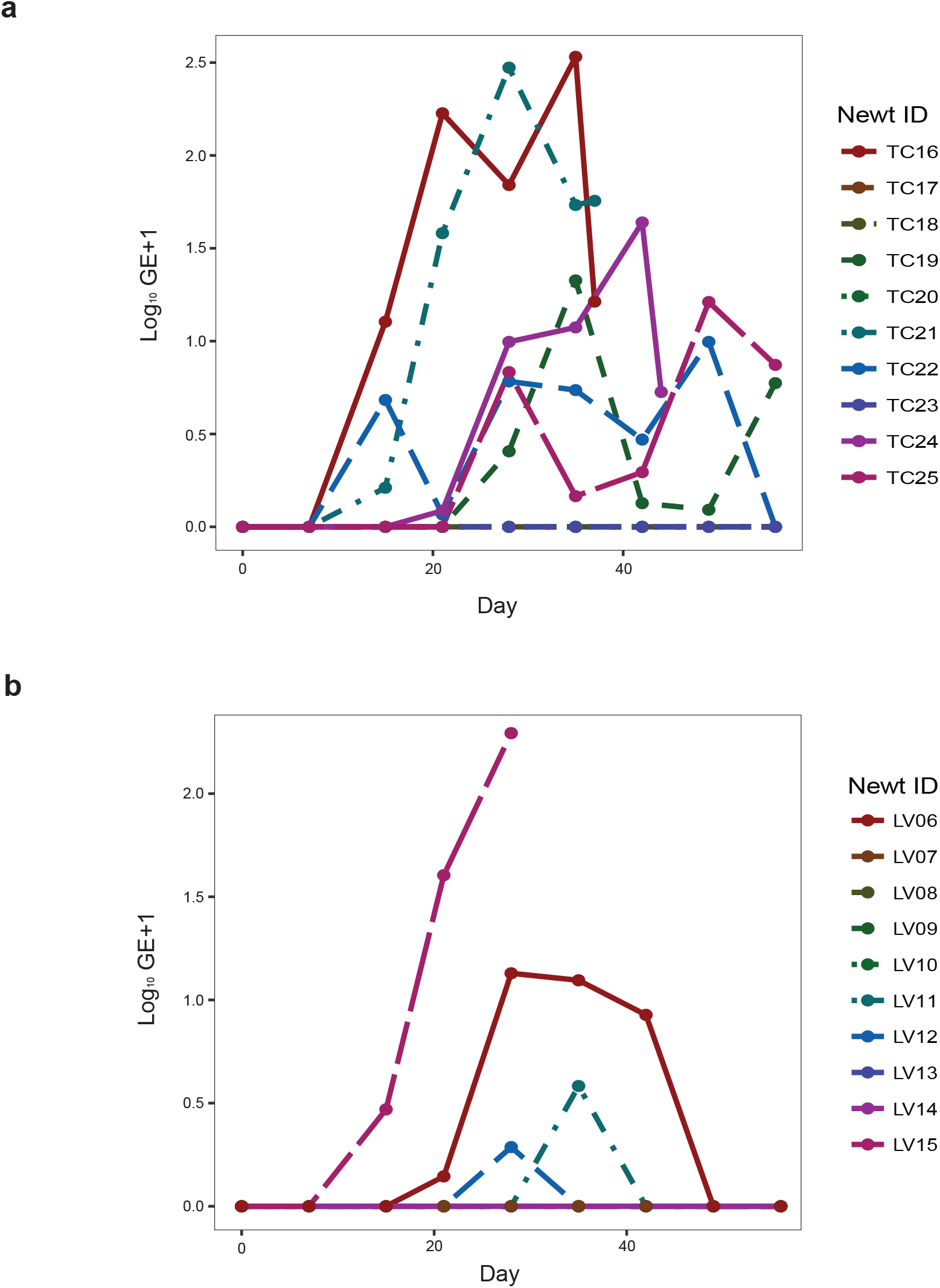
Plot of infection intensity for *Bsal* exposed animals for **(a)** *T. cristatus* **(b)** *L. vulgaris*.

Bacterial Shannon diversity differed based on day of sampling and mass for *T. cristatus* (day: F_(2,292)_=15.75, p=0.0001, mass: F_(1,13.7)_=4.68, *p*=0.05), but did not significantly alter over the course of the experiment in *L. vulgaris* (day: *p*=0.16, mass: *p*=0.96). Beta diversity differed only on day 28 based on treatment for *T. cristatus* (PERMANOVA Pseudo-F_(1,11)_=2.34, R^2^=0.108, *p*=0.045) and for both host species based on disease status (PERMANOVA *T. cristatus* Pseudo-F_(1,11)_=4.48, R^2^=0.199, *p*=0.002, *L. vulgaris* Pseudo-F_(1,11)_=4.95, R^2^=0.247, *p*=0.001, Fig. 4a, b) and survival (PERMANOVA *T. cristatus* Pseudo-F_(1,11)_=4.59, R^2^=0.204, *p*=0.006, *L. vulgaris* Pseudo-F_(1,11)_=2.44, R^2^=0.122, *p*=0.048). In *L. vulgaris*, on day 28 two individuals demonstrated microbiome perturbation, but cleared infection and returned to a microbiome state that did not differ significantly from control animals by day 56 (Fig. 4a). Indicator analysis of microbiome samples based on control, *Bsal* negative and *Bsal* positive animals from day 28 identified ten and six differentially abundant bacterial operational taxonomic units (OTUs) in *L. vulgaris* and *T. cristatus* respectively (Fig. 4c,d, SI Table 7). Both *L. vulgaris* and *T. cristatus* exhibited different indicator taxa profiles with the exception of *Stenotrophomonas* (Fig. 4e,f), which was strongly associated with *Bsal* infection in both species.

**Figure 4.**
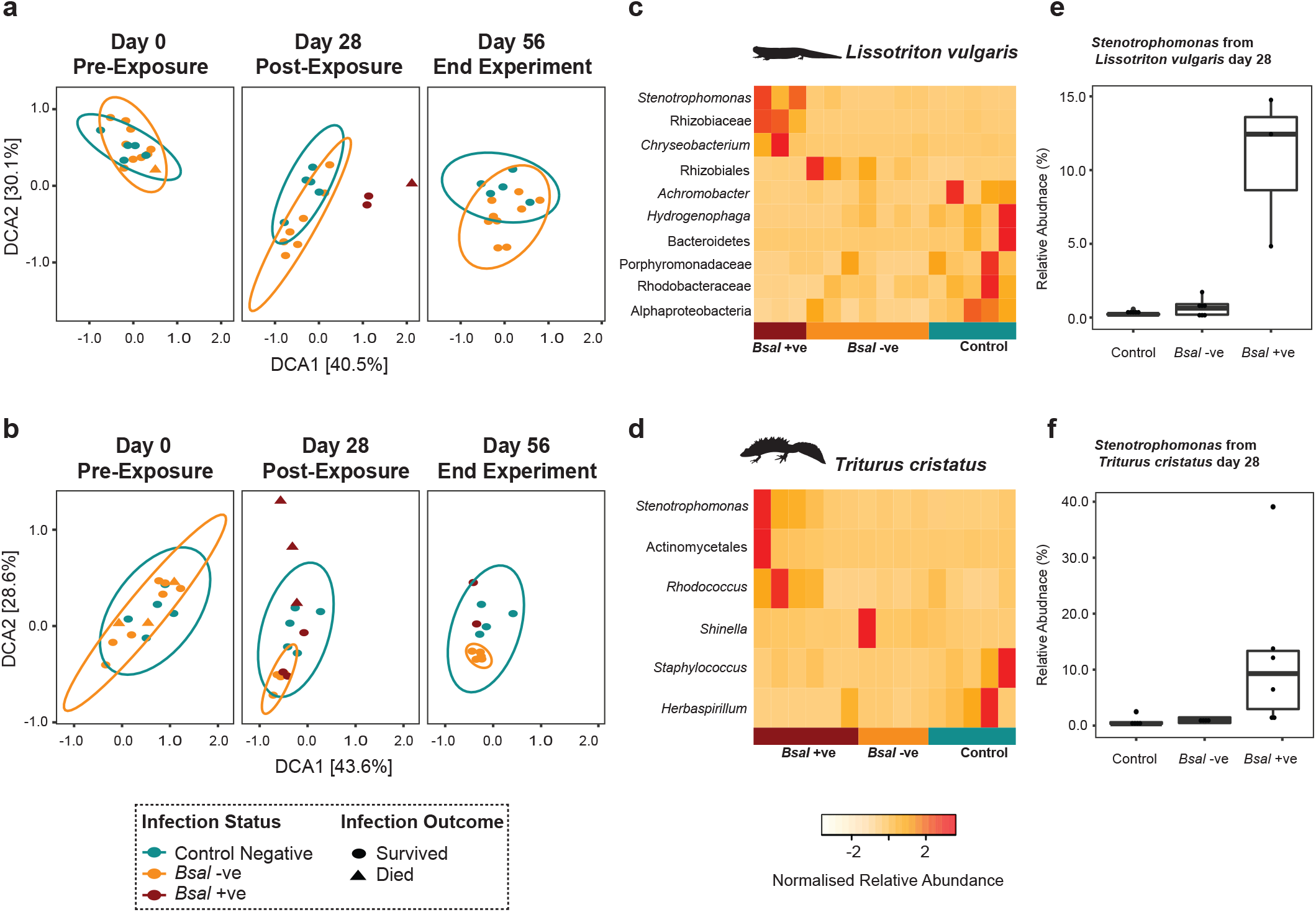
Detrended correspondence analysis (DCA) plots displaying temporal variation in bacterial beta diversity in **(a)** *L. vulgaris* **(b)** *T. cristatus*. Heatmap of normalised relative abundance of top bacterial indicator taxa associated with different disease outcomes in **(c)** *L. vulgaris* **(d)** *T. cristatus*. Boxplot of OTU016 *Stenotrophomonas* identified as an indicator OTU for *Bsal* infection in both **(e)** *L. vulgaris* and **(f)** *T. cristatus*.

## Discussion

We demonstrate that both captivity and exposure to *Bsal* impact the amphibian skin microbiome. The decrease in microbial Shannon diversity and changes in beta diversity associated with captivity supports findings of previous studies (Becker *et al*. 2014; Kueneman *et al*. 2016) and is likely correlated with a reduction in environmental microbes and changes to the host environment (Harrison *et al*. 2017). Importantly, both fungi and bacteria exhibited similar changes in alpha and beta diversity suggesting that commonalities exist in response to selection pressures across both microbial kingdoms. The perturbation in microbial diversity that occurs due to the transition from the wild to captivity may also change the ecological dynamics and functional capacity of the skin by altering host-microbe interactions in such a way that biases host- or microbe-mediated control of the microbiome (Foster *et al*. 2017). Specifically, the divergence in microbiome structure of different host species in captivity may be indicative of stronger host-mediated control, with host species differing in their selection of microbes in a common captive environment. This in turn may lead to decline of some potentially pathogenic taxa as signified by the reduction in OTUs belonging to the Chlamydiales (Fig. 2a,b) which have previously been associated with amphibian epizootics (Reed *et al*. 2000). Conversely, the reduced microbial diversity associated with captivity in both host species may also be a consequence of greater microbe-mediated control with diminished ecological resistance of the host skin microbiome rendering it more susceptible to potentially harmful invaders that may subsequently dominate the microbial community (Piovia-Scott *et al*. 2017). This hypothesis is supported by a reduction of putatively beneficial bacterial groups in captivity such as the Actinomycetales (Fig. 2a,b) which are well known for producing antimicrobial compounds (Berdy 2005). In addition, taxa belonging to the genus *Lysobacter* that have previously been associated with inhibition of other pathogenic chytrids (Brucker *et al*. 2008; Bates *et al*. 2018) showed a reduction in abundance in captivity. Captivity was also associated with an increase in abundance of the amphibian fungal pathogen *Basidiobolus ranarum* for both host species (though only significant in *T. cristatus*) further suggesting captive conditions may favour host-pathogen interactions as protective aspects of microbial diversity are lost. While the overall reduction in microbial diversity associated with captivity may impact host resistance to disease, the additional perturbation of microbial community structure may lead to dysbiotic effects (Zaneveld, McMinds & Thurber 2017). These findings suggest that captivity induced changes in the microbiome may reduce host capability to evade pathogens and potentially predispose such individuals towards adverse health outcomes. Importantly, this is the first study of its kind to measure the effect of captivity on fungal communities, which despite being overlooked relative to bacteria, have been shown in some cases to confer higher rates of chytrid inhibition (Kearns *et al*. 2017). In light of the major perturbation of fungal communities, it is imperative that future studies utilise a more holistic view of the microbiome to include kingdoms other than bacteria. Overall, our results highlight that host microbial ecology should carefully be considered when transferring animals from the wild to captivity to reduce possible microbiome disturbance, prevent proliferation of opportunistic pathogens and preserve microbes that are beneficial to host health. For *ex-situ* conservation programmes that aim to rescue or re-establish wild amphibian populations using captive-bred stock, the implications are complex and suggest that maintenance or ‘rewilding’ of the skin microbiome would be an essential aspect before reintroduction.

Exposure to *Bsal* resulted in mortality in both *L. vulgaris* and *T. cristatus*. While mortality was higher in *T. cristatus*, this was not statistically significant when compared to *L. vulgaris*. The intensity of infection was found to be higher in *T. cristatus*, which had a maximum GE of 338 compared to a maximum GE of 195 for *L. vulgaris*. These data, while preliminary, suggest that *T. cristatus* may be more susceptible to *Bsal* infection than *L. vulgaris*. Further, the greater zoospore burdens that *T. cristatus* harbour, coupled with their larger size and rates of zoospore shedding, suggest that they will be more important drivers of outbreak dynamics than *L. vulgaris*. Interestingly, our findings counter those of a previous study that found 100% mortality in *T. cristatus* and a species effect on survival (Martel *et al*. 2014). The increased survival of *T. cristatus* here compared to the prior study (Martel *et al*. 2014) may be explained by a range of factors. In particular, the studies used animals collected from different countries (the UK versus the Netherlands) and there will likely be differences in host genetics and immunity. In addition, the initial microbiome of the animals will almost certainly differ which may have an impact on the subsequent host microbial response to infection. These findings highlight the importance of investigating a geographically diverse range of hosts from the same species when conducting species level risks assessments of emerging pathogens. While this additional confirmation of *Bsal* induced death in *T. cristatus* warrants serious concern, the higher survivorship shown here may suggest a more optimistic outlook in the event of a wild *Bsal* outbreak for this species than previously predicted.

Bacterial community structure of individuals in the experiment differed significantly for both species based on infection status and survival. These findings are consistent with results of prior studies on *Bd* (Jani & Briggs 2014), suggesting that infection by *Bsal* and *Bd* are both associated with microbiome disruption. Two *L. vulgaris* individuals tested positive for *Bsal* and demonstrated significant microbiome perturbation that may be indicative of dysbiosis on day 28 of the experiment, but subsequently cleared infection by day 56. In both cases *Bsal* clearance was associated with a change in microbiome profile that was similar to that of control animals. Determining what underpins this recovery in *L. vulgaris*, whether driven by host factors such as secretion of antimicrobial peptides or *Bsal* inhibition by microbes, will be vital in determining the key parameters underpinning infection outcome. Indicator analysis revealed that the bacterial taxa associated with disease state were different based on host species with the exception of one OTU classified as *Stenotrophomonas*. Interestingly, a prior study demonstrated *Stenotrophomonas* as inhibitory to *Bsal* and several *Bd* genotypes (Muletz-Wolz *et al*. 2017). Meanwhile, another study identified an isolate of *Stenotrophomonas* as inhibitory to *Bd in-vitro*, however when applied as a probiotic on an amphibian host exposed to *Bd in-vivo*, mortality was higher than animals that were exposed to only *Bd* (Becker et al. 2015). Ultimately, determining whether *Stenotrophomonas* is associated with microbe mediated defence or synergistic growth with *Bsal* will require further experiments that take into account microbial function. While both newt species shared few taxa that were associated with disease outcome, the common perturbation in beta diversity associated with death in both *L. vulgaris* and *T. cristatus* may suggest that *Bsal* survival is not linked to specific taxa, but rather perturbation of a stable host microbiome resulting in an overall negative impact on health (Zaneveld, McMinds & Thurber 2017). Further support to this hypothesis is given by the return to a stable microbiome state in animals that cleared infection.

Our results build on the work of prior studies by demonstrating significant and potentially negative effects of captivity on the amphibian skin microbiome. In addition, we provide vital insight into the disease process of *Bsal* by demonstrating a close link between infection outcome and microbiome structure. Overall, our findings demonstrate that it is vital for host microbial ecology to be considered in future *Bsal* studies and captive based conservation programmes.

## Data accessibility

Sequence data have been deposited on the BioProject database under accession code PRJNA430498. All other data are available upon request from the authors.

## Acknowledgements

We thank A. Martel and F. Pasmans for providing the pathogen isolate used in the infection experiment. We are also grateful to Froglife for assistance with permits and field surveys. We thank P. Ghosh for help with experimental procedures and T. Garner for advice on experimental design. Finally, we wish to thank the Tedersoo group at the University of Tartu and the National Heart and Lung Institute for assistance with mycobiome sequencing. This research was funded by the Leverhulme Trust grant RPG-2014-273.

## Author contributions

K.A.B. and V.L.M. conducted field surveys and animal experiments. K.A.B., V.L.M., K.H., X.A.H., and J.M.G.S. performed data processing and analysis. K.A.B., M.C.F., S.P., X.A.H., V.L.M. and J.M.G.S. wrote the manuscript.

## Ethic statement

Animal work was conducted under UK Home Office Project Licence PPL 70/8402 held by Matthew Fisher and was reviewed by the Imperial College London Animal Welfare Ethical Review Board for approval. The experiment was carried out in accordance with The Animals (Scientific Procedures) Act of 1986 Directive 2010/63/EU and followed all of the Codes of Practice which reinforce this law, including all elements of housing, care and euthanasia. Animals were collected in the wild under Natural England Licence 2015-15771-SCI-SCI and following ethical review by the board of Froglife.

